# The processing of stress in a foreign language modulates functional antagonism between default mode and attention network regions

**DOI:** 10.1101/2022.12.21.521156

**Authors:** Lars Rogenmoser, Michael Mouthon, Faustine Etter, Julie Kamber, Jean-Marie Annoni, Sandra Schwab

## Abstract

Lexical stress is an essential element of prosody. Mastering this prosodic feature is challenging, especially in a stress-free foreign language for individuals native to a stress-fixed language, a phenomenon referred to as stress deafness. By using functional magnetic resonance imaging, we elucidated the neuronal underpinnings of stress processing in a stress-free foreign language, and determined the underlying mechanism of stress deafness. Here, we contrasted behavioral and hemodynamic responses revealed by native speakers of a stress-free (German; *N* = 38) and a stress-fixed (French; *N* = 47) language while discriminating pairs of words in a stress-free foreign language (Spanish). Consistent with the stress deafness phenomenon, French speakers performed worse than German speakers in discriminating Spanish words based on cues of stress but not of vowel. Whole-brain analyses revealed widespread bilateral networks (cerebral regions including frontal, temporal and parietal areas as well as insular, subcortical and cerebellar structures), overlapping with the ones previously associated with stress processing within native languages. Moreover, our results provide evidence that the structures pertaining to a right-lateralized attention system (i.e., middle frontal gyrus, anterior insula) and the Default Mode Network modulate stress processing as a function of the proficiency level. In comparison to the German speakers, the French speakers activated the attention system and deactivated the Default Mode Network to a stronger degree, reflecting attentive engagement, likely a compensatory mechanism underlying the “stress-deaf” brain. The mechanism modulating stress processing argues for a rightward lateralization, indeed overlapping with the location covered by the dorsal stream but remaining unspecific to speech.

## 1. Introduction

Mastering the prosodic aspect of a language relies considerably on the correct processing of lexical stress (abbr. stress), which is characterized by the differential accentuation of a particular syllable within words. Stress processing (SP) is cognitively demanding (Dupoux et al., 2001), especially in a foreign language, whereby its difficulty depends on the native/primary language (L1) of the individual itself and whether the L1 is stress-*free* or stress-*fixed* (Dupoux et al., 1997). In stress-free languages (such as English, German, Spanish), the position of the stressed (or salient) syllable varies amongst words, sometimes with a contrastive and semantically determining function. Thus, the proper acquisition of a stress-free language requires the additional effort of associating every single word with specific stress patterns. In stress-fixed languages (such as French, Hungarian), by contrast, the stressed syllable within words is invariant and does not bare any distinctive properties. Due to this inherent nature, stress-fixed L1 speakers do not sensitize to stress features, disadvantaging them over stress-free L1 speakers when performing stress-related tasks, such as discriminating between words with differential stress patterns. This effect is referred to as stress deafness (SD) (Dupoux et al., 2008, 2001, 1997), a robust finding meanwhile documented in many behavioral studies (Alfano et al., 2010; Dupoux et al., 2001, 1997; Honbolygó et al., 2019; Peperkamp et al., 2010; Schwab and Dellwo, 2017). These studies have ascribed SD to various cognitive factors mainly underlying working memory (Dupoux et al., 2008, 2001, 1997; Heisterueber et al., 2014; Honbolygó et al., 2019), a conclusion also supported by brain response recordings using electroencephalography during a series of stress-related detection tasks (Lu et al., 2018; Schwab et al., 2020). So far, imaging studies have associated SP with the involvement of brain regions, mainly overlapping with the two speech-specific pathways (ventral and dorsal stream; (Bornkessel-Schlesewsky et al., 2015; Bornkessel-Schlesewsky and Schlesewsky, 2013; DeWitt and Rauschecker, 2012; Sammler et al., 2015), particularly with temporal (superior temporal gyrus and sulcus, STG/S; Aleman et al., 2005; Klein et al., 2011; Domahs et al., 2013; Heisterueber et al., 2014; Honbolygó et al., 2020), frontal (inferior and middle frontal gyrus, I/MFG; premotor; supplementary motor area, SMA; Aleman et al., 2005; Klein et al., 2011; Domahs et al., 2013; Heisterueber et al., 2014; Kandylaki et al., 2017) and parietal structures (angular gyrus, AnG; precuneus, PCu; Klein et al., 2011; Domahs et al., 2013; Heisterueber et al., 2014; Kandylaki et al., 2017). Some of the studies have further concluded the involvement of the insula (Aleman et al., 2005; Heisterueber et al., 2014; Klein et al., 2011), basal ganglia and cerebellum (Heisterueber et al., 2014; Klein et al., 2011). However, these insights were gained from studies on participants limited within one L1, where mainly laboratory stimuli (e.g., pseudowords) were used to ensure certain stress-related responses up to mimicking SD.

The aim here is to elucidate the brain while realistically processing stress in a stress-free foreign language (i.e., Spanish), and to determine the neuronal underpinnings of actual SD by contrasting the responses revealed by speakers of a stress-free (i.e., German) and a stress-fixed (i.e., French) L1. In the present study, samples of both German and French speakers underwent functional magnetic resonance imaging (fMRI) while discriminating well-controlled trisyllabic Spanish word pairs, systematically compiled based on a specific triplet-approach (Dupoux et al., 1997). The participants were instructed to judge as quickly and accurately as possible whether or not the two words of pairs differed. Word pairs differed either in terms of the stress or vowel information (Klein et al., 2011). Given that stress-fixed speakers do not encode stress, we hypothesize French speakers to underperform only in the stress condition over German speakers, who may be able to exploit a certain transfer effect due to stress-encoding strategies acquired in a stress-free L1. Due to growing evidence for general cognitive factors considerably accounting for SD (Alfano et al., 2010; Dupoux et al., 2008, 2001, 1997; Honbolygó et al., 2019; Lu et al., 2018; Peperkamp et al., 2010; Schwab et al., 2020; Schwab and Dellwo, 2017), we hypothesize SD to underlie an altered mode of speech-independent brain processing.

## 2. Materials and Methods

### 2.1 Participants

Ninety-one right-handed students enrolled in this study. Six participants were removed due to excessive head movements or technical issue during data acquisition. Eighty-five remained for the data analysis, with 47 native French speakers (age: 23.06 ± 2.39 years; 28 females) and 38 native German speakers (age: 22.74 ± 2.34 years; 27 females). The two samples were comparable regarding age (*t*_83_ = 0.63, *P* = 0.53) and the distribution of the sexes (*Χ*^*2*^_1_ = 1.21, *P* = 0.27). The participants had no knowledge of any stress-free romance languages including Spanish, Italian, or Portuguese. All participants gave written informed consent, and were paid for their participation. The work has been carried out in accordance with The Code of Ethics of the World Medical Association (Declaration of Helsinki) for experiments involving humans, and the study was approved by the Ethics Committee of the Psychology Department of the University of Fribourg.

### 2.2 Experimental Design

All participants performed a discrimination task during the fMRI session. In this task, the participants were exposed to a stream of Spanish word pairs, presented binaurally via MR Confon system (Magdeburg, Germany). The participants were instructed to judge as quickly and accurately as possible whether or not the two words of the presenting pairs differed by pressing right-handedly on one of two response buttons. The words differed in terms of two violation conditions, namely regarding the stress and the vowel information. Both conditions were presented in blocks, which were alternated across the session. In total, 32 blocks were presented in a pseudorandomized manner. The duration of one block was of 22.2 s, followed by a blank screen with a duration of 8 s. Each block consisted of 6 trials containing word pairs, whereby in half of the trials the words differed and in the other half they did not. Each block was initiated visually via LCD monitor (NordicNeuroLab, Bergen, Norway) with the instruction and announcement of the upcoming condition, which lasted 1 s. Afterwards, a fixation cross with a duration of another 1 s was presented, followed by the particular trials. The trials were 1.7 s in duration, with the two words separated by an interval of 500 ms and each word containing a length of 600 ms. The inter-trial interval was 2 s. The task took approximately 18 min and was divided into two equal scan runs with a 1-min break in between. Before scanning, the participants were given 12 practice trials to get familiar with the task. Stimulus presentation as well as behavior collection was controlled by the software E-Prime 3.0 (Psychology Tools, Inc., Pittsburgh, PA, USA).

### 2.3 Stimulus Material

Twenty-four trisyllabic Spanish verbs were compiled for the task (for word list, see Table 1). Each verb appeared in three inflectional forms; (i) the first form ending with -o together with the stress on the penultimate syllable (first person, singular present indicative), (ii) the second form segmentally identical to the first one also ending with -o, but with the stress on the final syllable (third person, singular simple past indicative), (iii) and the third form segmentally different from the first, instead ending with -e but with stress on the penultimate syllable (first/third person, singular present subjunctive). Within each triplet, pairs of the first form together with the second functioned as stress condition and pairs of the first together with the third functioned as vowel condition. Per triplet, pairs with no manipulation were additionally added. These pairs contained mere duplicates of either the second (stress condition) or the third form (vowel condition). In total, a pool of 96 pairs was generated (i.e., 24 verbs x 2 conditions x 2 same/different). Finally, the number of pairs was doubled with a second stimulus set of the same pool only with inverted presentation order of the words within the pairs, resulting in 192 pairs in total. Each word was digitally recorded, uttered by a female native speaker of Castilian Spanish. Later, the recorded words were trimmed into lengths of 600 ms using the software Praat (Boersma and Weenink, 2011).

**Table 1.**
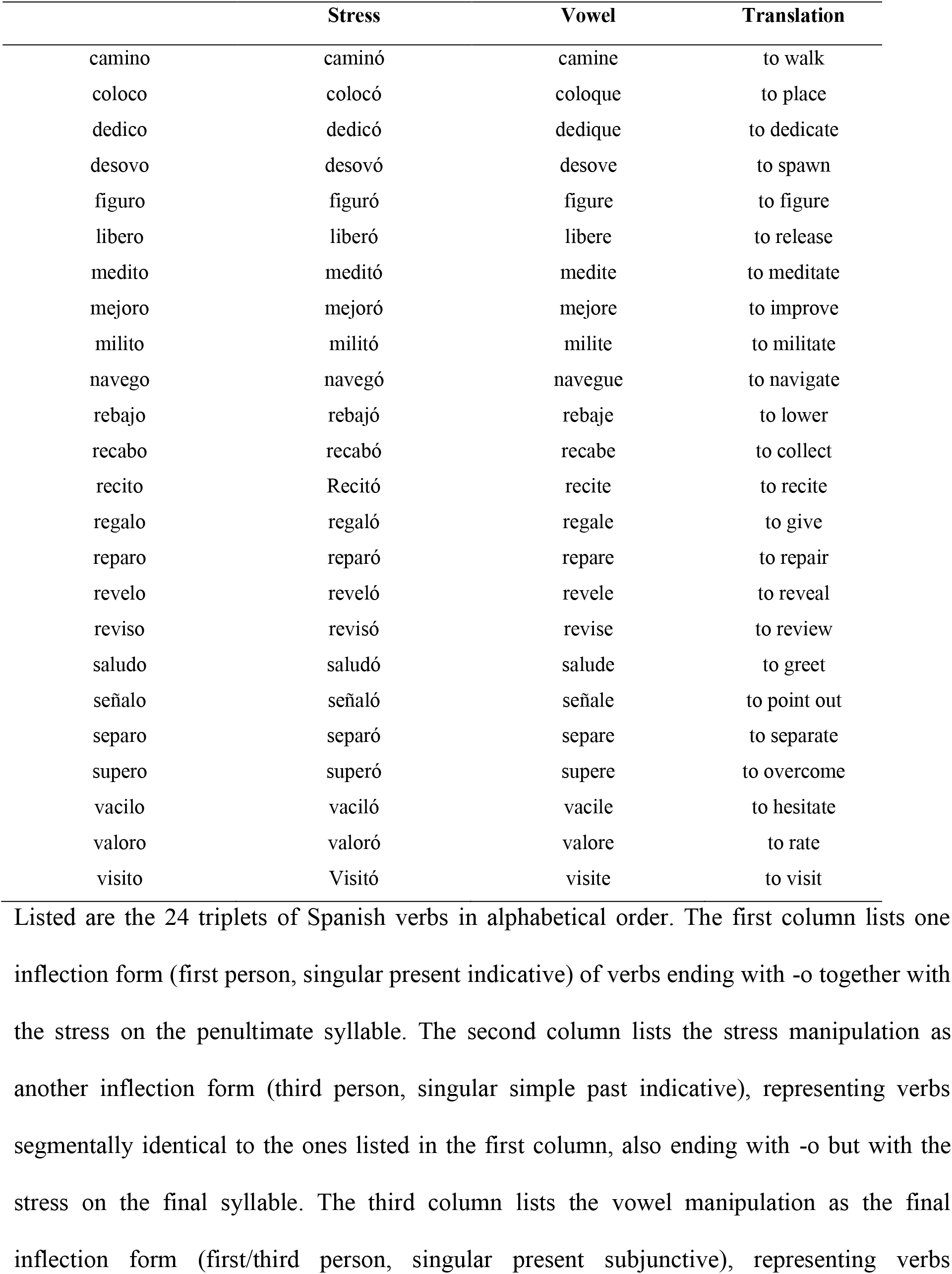
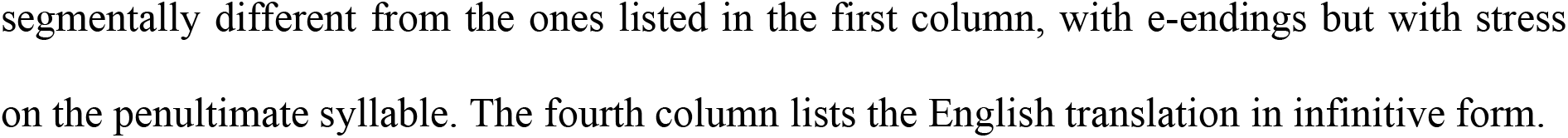
Word list.

### 2.4 Image Acquisition and Preprocessing

Images were acquired using a 3 tesla MRI scanner (Discovery MR750; GE Healthcare, Waukesha, Wisconsin) equipped with a 32-channel head coil. For each participant, a high-resolution T1-weighted anatomical scan of the brain was acquired from the coronal plane with 270 slices and a voxel size of 0.86x 0.86 × 1 mm (FOV = 22 cm, acquisition matrix: 256 × 256, TR = 7.3 ms, TE = 2.8 ms, Flip angle = 9°, Prep time = 900 ms, parallel imaging acceleration factor (PIAF) = 1.5). Blood oxygen level-dependent (BOLD) functional images were acquired with a T2*-sensitive echo-planar image (EPI) gradient-echo pulse sequence. The BOLD was used as an index of local increases in brain activity (Kwong et al., 1992). Each functional run consisted of 264 dynamic volumes with axial contiguous ascending acquisitions and a voxel size of 2.3 × 2.3 × 3 mm (FOV = 22 cm, acquired matrix: 96 × 96, number of slices = 36, slice thickness = 3 mm, inter-slices space = 0.5 mm, TR = 2000 ms, TE = 30 ms, Flip angle = 85°, PIAF=2). To assure a steady-state magnetization of the tissue, each run started with a dummy scan of 8 s. After the experimental task, a fieldmap was acquired in order to correct for the distortion of the static magnetic field during post-processing. This required two FAST SPGR sequences with distinct Echo Time and the same space coverage as the functional EPI (TR = 50 ms, TE1 = 4.9 ms, TE2 = 7.3 ms, flip angle = 45°). MRI data preprocessing and analyses were conducted with the Statistical Parametric Mapping Software (SPM12) running on MATLAB R2016b (MathWorks, MA). The following pipeline was used to preprocess the functional images: setting the origin on the anterior commissure, slice timing, computation of the Voxel Displacement map (VDM) (using the FieldMap2.1 toolbox (Andersson et al., 2001)), spatial realignment and unwarping (using VDM previously created), normalization into the MNI coordinate system with a voxel size of 3x3x3 mm based on the Unified-Segmentation procedure of the co-registered T1-weighted anatomical image to fMRI images, and smoothing with a Gaussian kernel of 8-mm full-width-at-half-maximum (FWHM). The ArtRepair toolbox was used to detect volumes with fast motion (>0.5 mm/TR) during fMRI recording. A general linear model (GLM) was used to analyze the resulting preprocessed images at the individual subject level. The fMRI signal was modeled as condition-specific block of 22.63 s of duration convolved with the hemodynamic response function (HRF). To remove low frequency noise and signal drifts, a high-pass filter with a 1/128 Hz threshold was applied at time series from each voxel. An autoregressive function (AR(1)) was implemented to correct for temporal correlations between neighboring voxels in the whole brain. The simple positive contrasts for each experimental condition and each participant were estimated (fixed-effect) and subjected to the group analysis (random effect).

### 2.5 Statistical Analysis

Regarding behavioral data, the mean RT and accuracy scores obtained from each condition by each participant were imported into the SPSS software for inferential statistics. The data were subjected to two-way mixed analyses of variance (ANOVA) with L1 (fixed, free) as between-factor and task condition (stress, vowel) as within-factor. Statistical analyses were adjusted for non-sphericity using Greenhouse-Geiser Epsilon when equal variances could not be assumed. Significant interaction effects were further inspected using post-hoc pairwise two-tailed *t*-tests. The Bonferroni procedure was applied to correct for multiple comparisons.

The fMRI data were subjected to a two-way mixed factorial ANOVA with L1 as between-factor (fixed, free) and task condition as within-factor (stress, vowel) using the Sandwich Estimator Toolbox (Guillaume et al., 2014). For each group, contrasts (stress > vowel, vowel > stress) were subjected to non-parametric one-sample *t*-tests. The results were studied on the whole brain space with the statistical threshold FWE corrected for multiple comparison at the peak level (*P*_FWE_ < 0.05) with the Threshold-Free Cluster Enhancement Estimator (TFCE) (Smith and Nichols, 2009). Anatomical locations were checked with the neuromorphometrics probabilistic atlas provided by SPM12. All the coordinates derived from these analyses are given in the MNI space.

## 3. Results

### 3.1 Behavioral results

The reaction time (RT) and accuracy scores achieved in discriminating the stress-free foreign word pairs during the fMRI session are depicted in Fig. 1. Two-way mixed analyses of variance (ANOVA), with condition (stress, vowel) as within- and L1 (stress-fixed/French, stress-free/German) as between-factors, revealed that the condition had no impact on the RT (*F*_1, 83_ = 0.73, *P* = 0.395, *η*^*2*^*p* = 0.009) but had one on the accuracy rate (*F*_1, 83_ = 107.35, *P* = 1.27 × 10^−16^, *η*^*2*^*p* = 0.564), with stress being the more challenging condition. Generally, the French speakers responded slower (*F*_1, 83_ = 4.67, *P* = 0.033, *η*^*2*^*p* = 0.053) and made more errors (*F*_1, 83_ = 5.89, *P* = 0.017, *η*^*2*^*p* = 0.066). This performance drop was especially pronounced in the stress condition, as revealed by the between-within interaction effects (RT: *F*_1, 83_ = 17.05, *P* = 8.6 × 10^−5^, *η*^*2*^*p* = 0.17; accuracy: *F*_1, 83_ = 28.75, *P* = 7.27 × 10^−7^, *η*^*2*^*p* = 0.257), confirming a SD effect among the French speakers. Post-hoc pairwise comparisons (Bonferroni-adjusted at the level of α = 0.05 for 4 two-sided *t*-tests) corroborated that the French speakers responded slower (*t*_66.32_ = 3, *P* = 0.016, *d* = 0.68) and with less precision only in the stress condition (*t*_83_ = 4.25, *P* = 2.44 × 10^−4^, *d* = 0.92), whereas L1 differences in the vowel condition did not reach any significance (RT: *t*_65.42_ = 1.25, *P* = 1.13, *d* = 0.25; accuracy: *t*_46.29_ = 1.95, *P* = 0.228, *d* = 0.46). Furthermore, the post hoc tests revealed that both groups discriminated words based on the vowel information with more precision (German: *t*_37_= 3.89, *P* = 0.002, *d* = 0.63; French: *t*_46_= 10.73, *P* = 1.64 × 10^−14^, *d* = 1.57) and that the stress-specific delaying effect was limited to the French speakers’ responses (French: *t*_46_= 3.38, *P* = 0.006, *d* = 0.49; German: *t*_37_= 2.58, *P* = 0.056, *d* = 0.42).

**Figure 1.**
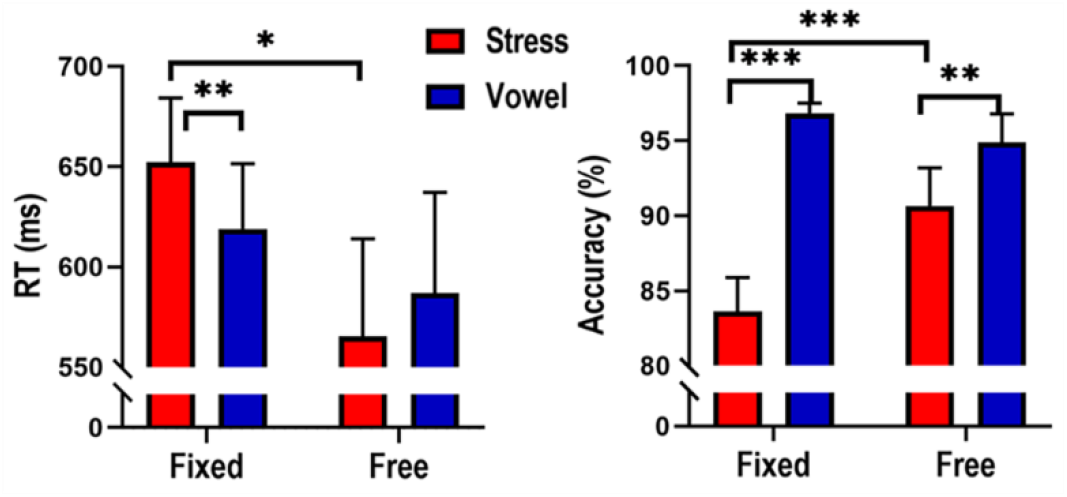
Performance in discriminating pairs of words in a stress-free foreign language (Spanish). The mean reaction time (RT, left) as well as accuracy scores (right) are depicted across both conditions (stress, vowel) for both native languages (stress-fixed/French *N* = 47, stress-free/German *N* = 38). Stress “deafness” is indicated by the poorer performance among stress-fixed (French) participants in the stress condition. The bars depict 95% confidence intervals. Two-tailed Bonferroni-adjusted \*\*\**P* < .001, \*\**P* < .01 \**P* < .05.

### 3.2 Imaging results

A two-way mixed ANOVA, with L1 (stress-fixed/French, stress-free/German) as between- and condition (stress, vowel) as within-factors, revealed no significant effect of group on the brain activation, but one of condition and one of between-within interaction (*P* < 0.05 with *α*-error corrected family-wisely at voxel level, FWE; threshold-freely clustered). The main effect of condition comprised 3 significant clusters of activation, peaking in the right opercular inferior frontal gyrus (OpIFG), left cerebellum exterior (CbE), and right supramarginal gyrus (SMG). The cluster with the activation peaking in the OpIFG (*k* = 24700 voxels) extended across the bilateral frontal region (i.e., IFG, MFG, SMA) and included temporal and occipital areas as well as structures within the cingulate gyrus and insula. Table 2 details the 3 clusters of the main effect of condition, and Fig. 2 illustrates their distribution of activation mapped onto a standard (Montreal Neurological Institute, MNI) brain template. The between-within interaction effect comprised 6 significant clusters of activation, peaking in the left posterior cingulate gyrus (PCG), left medial superior frontal gyrus (MSFG), left parahippocampal gyrus (PHG), right AIns and in two spots within the right MFG. The cluster surrounding the PCG (*k* = 2306 voxels) included bilaterally the middle occipital gyri (MOG) together with the PCu as well as the right AnG.

**Table 2.**
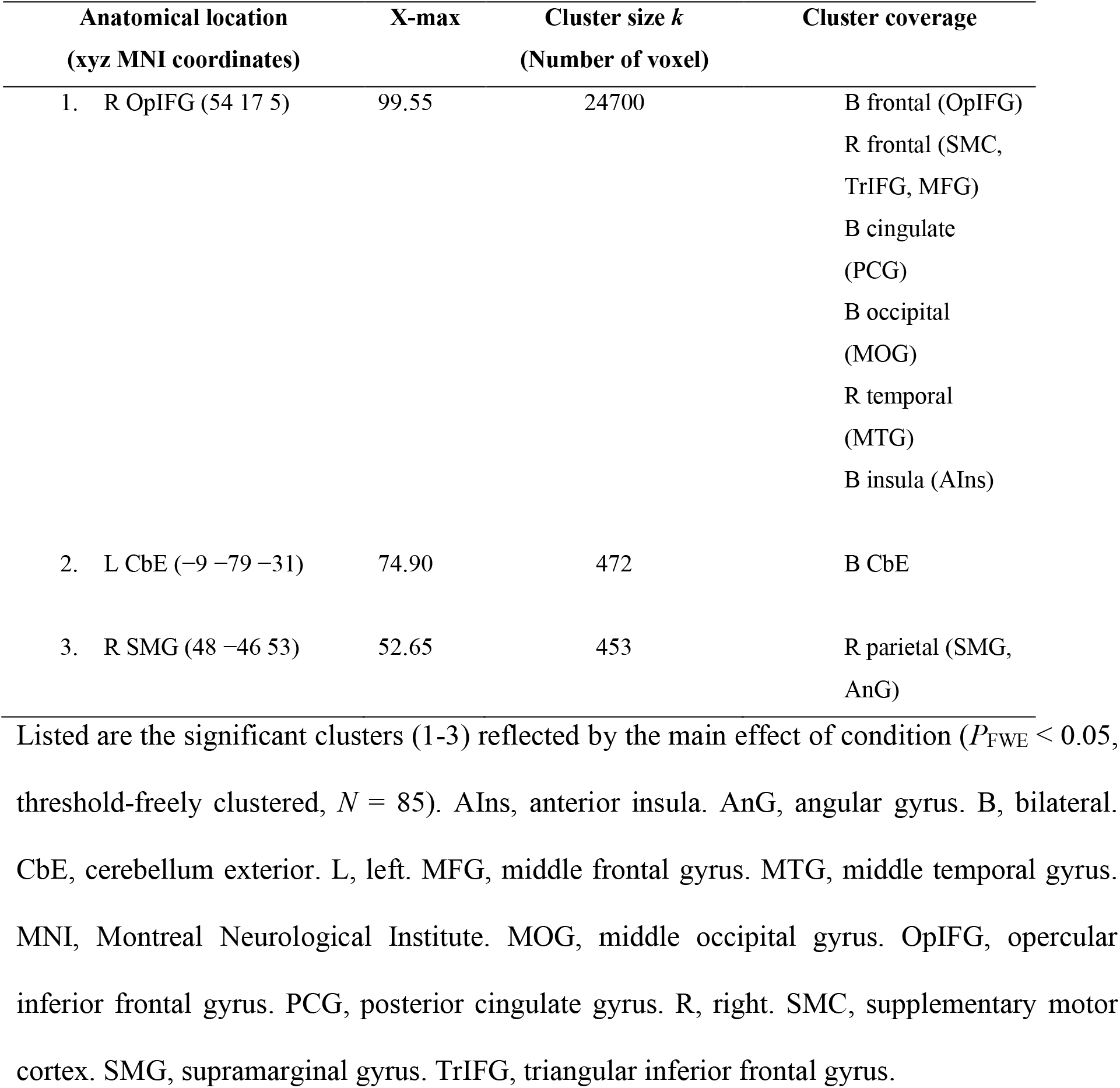
Main effect: Brain regions involved as a function of the processing condition.

**Figure 2.**
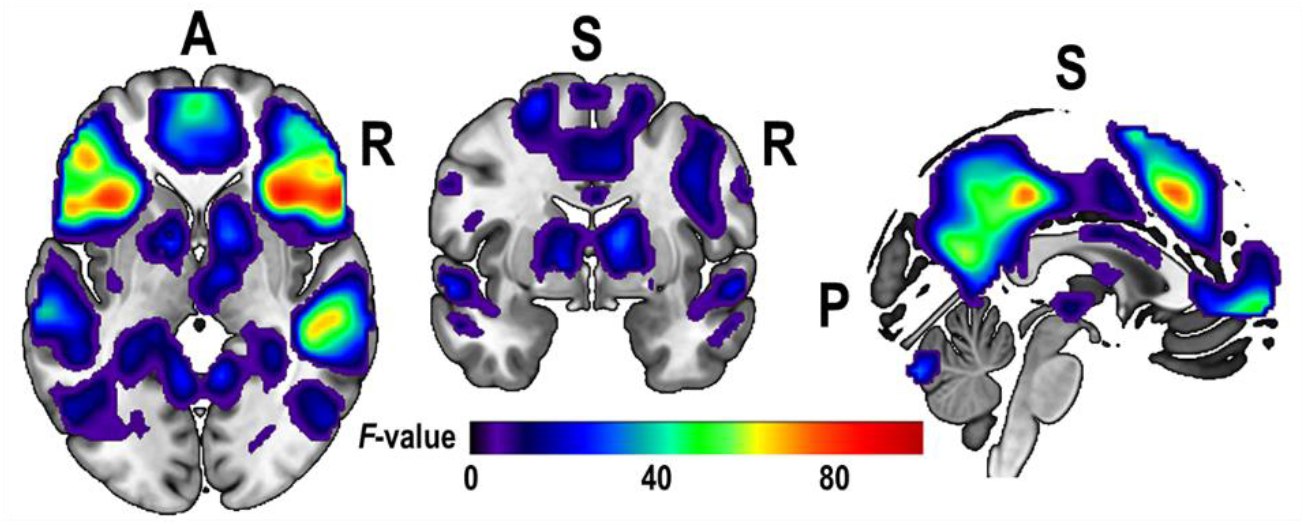
Main effect of the processing condition. Activation (*P*_FWE_ < 0.05, threshold-freely clustered, *N* = 85) is mapped onto a standard brain template (MNI152) with the xyz-coordinates set at zero. For cluster details, see Table 2. Apparent are the axial (left), coronal (middle) and sagittal (right) view. A, anterior. P, posterior. R, right. S, superior.

Table 3 details the 6 clusters of the interaction effect, and Fig. 3 illustrates their distribution of activation mapped onto an MNI brain template. The interaction effect was due to the French speakers exhibiting a differentiated task-dependent activation response across all clusters. In comparison to the German speakers, the French speakers showed an increase in activity when processing stress over vowel specifically in the clusters located around attention-related brain structures (i.e., right MFG and AIns (Fox et al., 2006; Japee et al., 2015; Sridharan et al., 2008; Uddin, 2015)). In the remaining clusters that covered brain structures pertaining to the Default Mode Network (DMN; i.e., PCG, MSFG, PHG (Buckner et al., 2008; Fox et al., 2005; Raichle, 2015; Ward et al., 2014)), the French speakers exhibited an inverted response pattern, namely a deactivation when processing stress over vowel (or an activation when processing vowel over stress). Fig. 4 illustrates this interaction effect plotted across the 6 clusters, pertaining to both an attention-related system and the DMN.

**Table 3.** Interaction effect: Brain regions involved as a function of the processing condition and the native language.

| Anatomical location<br>(xyz MNI coordinates) | X-max | Cluster size $k$<br>(Number of voxel) | Cluster coverage |
| --- | --- | --- | --- |
| 1. L PCG (−3 −34 44) | 25.5 | 2306 | B cingulate (PCG)<br>B occipital (MOG)<br>B parietal (PCu)<br>B parietal (AnG) |
| 2. L MSFG (−3 59 −1) | 19.28 | 161 | B MSFG |
| 3. R MFG (42 23 26) | 17.83 | 204 | R MFG |
| 4. R MFG (36 56 8) | 16.4 | 16 | R MFG |
| 5. L PHG (−30 −34 −13) | 17.48 | 54 | L PHG |
| 6. R AIns (33 20 −4) | 17.21 | 32 | R Ains |

Listed are the significant clusters (1-6) reflected by the interaction effect ( $P_{\text{FWE}} < 0.05$ , threshold-freely clustered; stress-fixed, French $N = 47$ ; stress-free German $N = 38$ ). AIns, anterior insula. AnG, angular gyrus. B, bilateral. L, left. MFG, middle frontal gyrus. MSFG, medial superior frontal gyrus. MNI, Montreal Neurological Institute. MOG, middle occipital gyrus. PCG, posterior cingulate gyrus. PCu, precuneus. PHG, parahippocampal gyrus. R, right.

**Figure 3.**
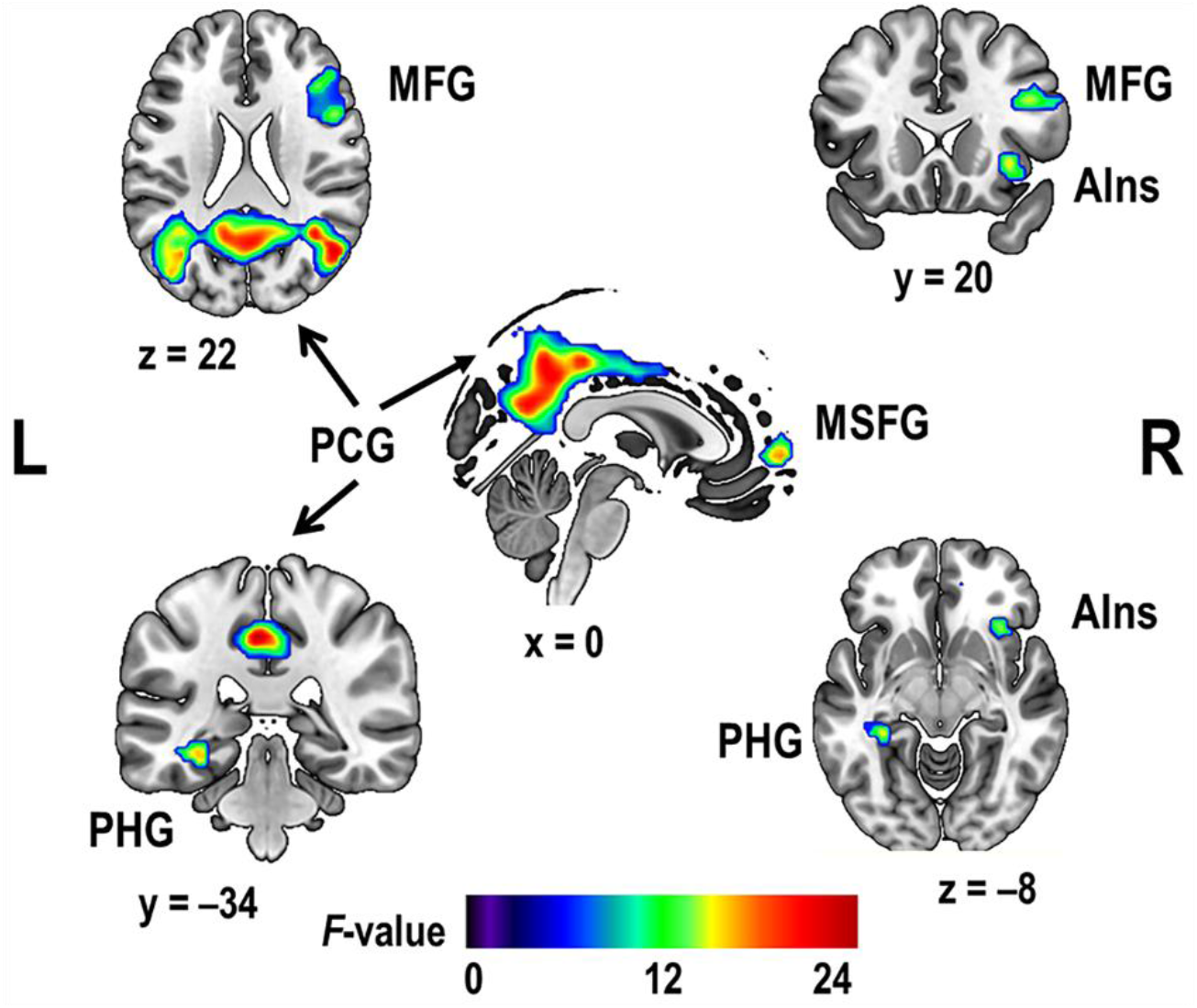
Interaction effect as a function of the processing condition and native language. Activation (*P*_FWE_ < 0.05, threshold-freely clustered, stress-fixed/French *N* = 47, stress-free/German *N* = 38) is mapped onto a standard brain template (MNI152). For cluster details, see Table 3. AIns, anterior insula. L, left. MFG, middle frontal gyrus. MSFG, medial superior frontal gyrus. PCG, posterior cingulate gyrus. PHG, parahippocampal gyrus. R, right.

**Figure 4.**
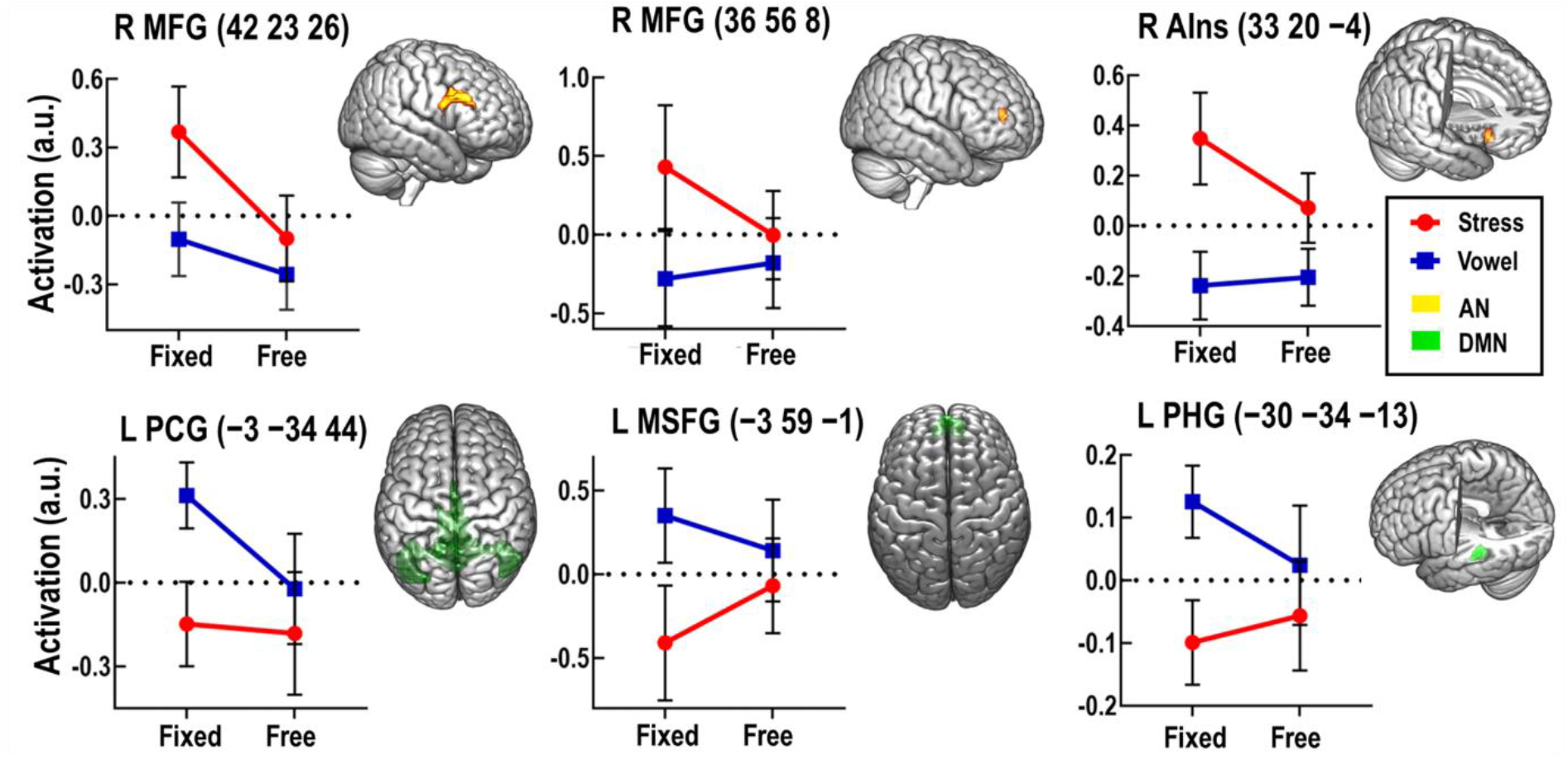
Mean activation values of the interaction effect as a function of the processing condition and native language. The values are plotted across the 6 significant clusters (*P*_FEW_ < 0.05, threshold-freely clustered, stress-fixed/French *N* = 47, stress-free/German *N* = 38), pertaining to the attention network (AN, top) and the Default Mode Network (DMN, bottom). For cluster details, see Table 3. Montreal Neurological Institute coordinates (xyz) are indicated in brackets. AIns, anterior insula. L, left. MFG, middle frontal gyrus. MSFG, medial superior frontal gyrus. PCG, posterior cingulate gyrus. PHG, parahippocampal gyrus. R, right. The bars depict 95% confidence intervals.

To identify the differential brain regions contributing to the processing of stress and vowel, respectively, we contrasted the underlying activation patterns revealed from both processing conditions, with either stress against the vowel processing (i.e., stress > vowel, stress-contrast) or vice versa (vowel > stress, vowel-contrast). For each group, we subjected these contrasts to non-parametric one-sample *t*-tests. The separate contrasts revealed significant activations at regions (*P*_FWE_ < 0.05, threshold-freely clustered) overlapping largely across both groups, although the particular clusters differed in terms of number, peak location, coverage and size. Details of the clusters are listed in Table 4 (for the French) and Table 5 (for the German). Fig. 5 illustrates both groups’ distribution of the activation mapped onto an MNI brain template. In both groups, the stress-contrast revealed activation over bilateral fronto-temporal areas, the right AIns, thalamus, left CbE and pallidum. Among the French speakers, the *t*-test revealed further activation in the right inferior parietal lobule (i.e., SMG, AnG). The vowel-contrast revealed, in both groups, activation over the bilateral parieto-occipital and cingulate areas together with the MFG. Finally, the *t*-tests identified rather smaller clusters covering the right PHG and left thalamus (*k* = 2 voxel) among the German speakers, and within the right posterior insula (*k* = 7 voxel) among the French speakers.

**Table 4.**
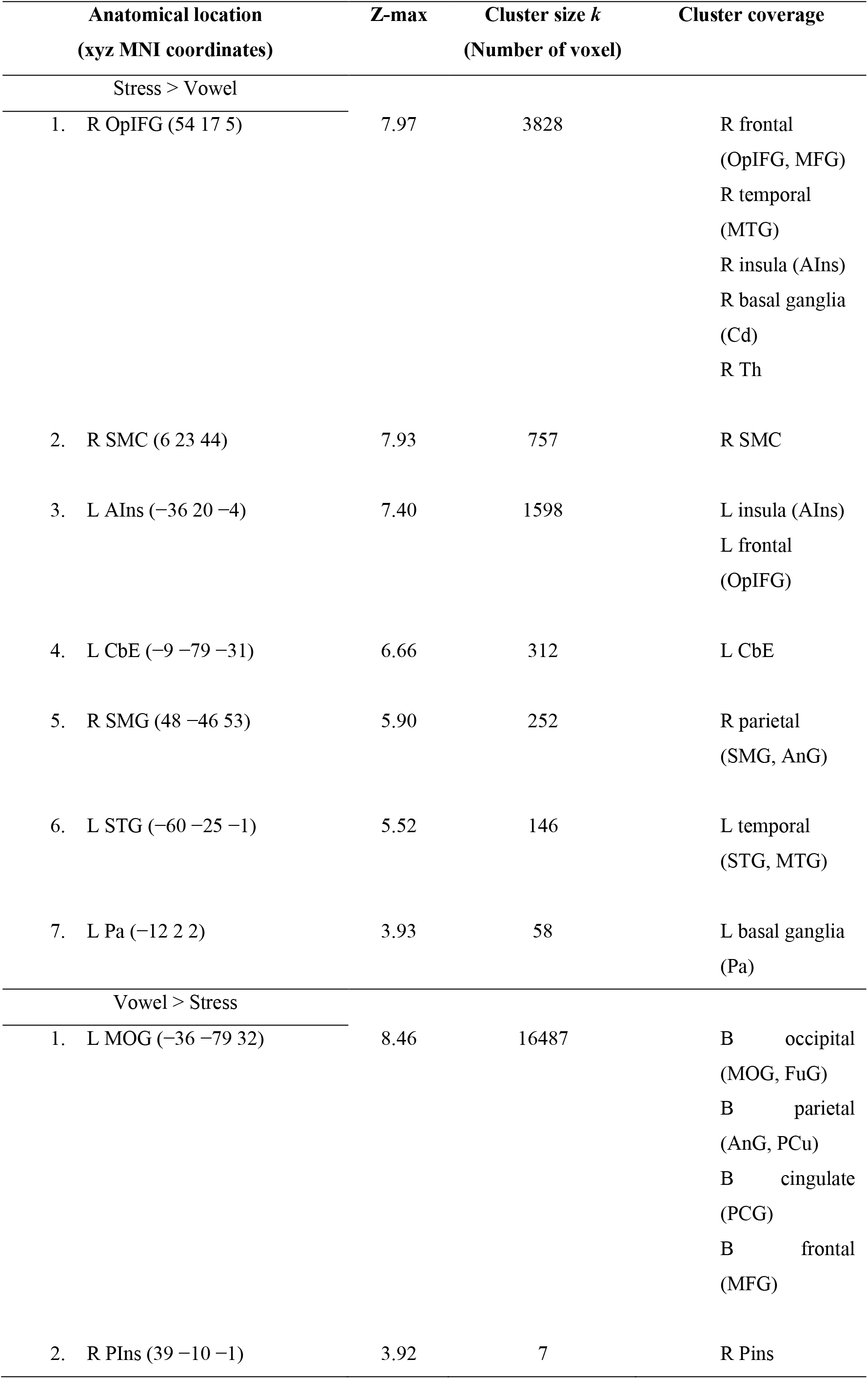
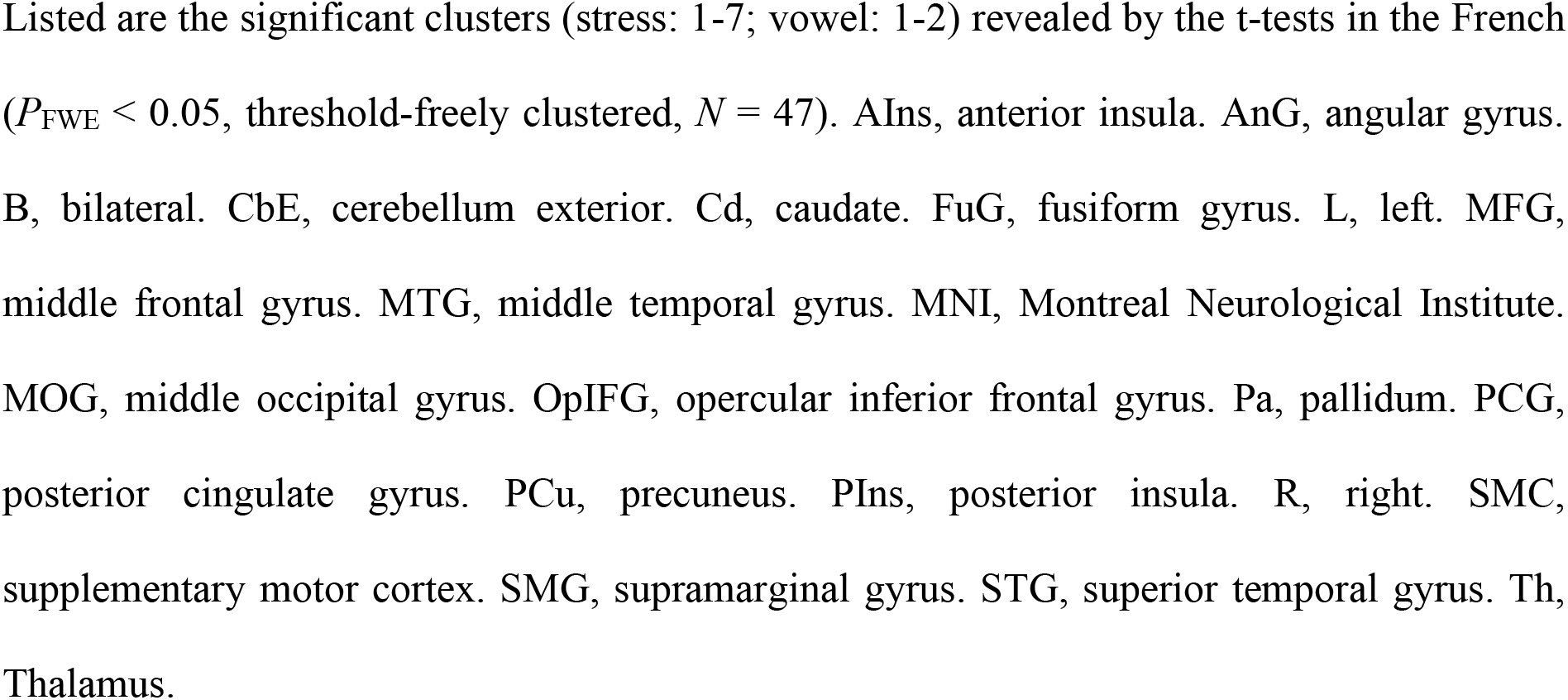
French speakers: Brain regions involved in stress and vowel processing.

**Table 5.**
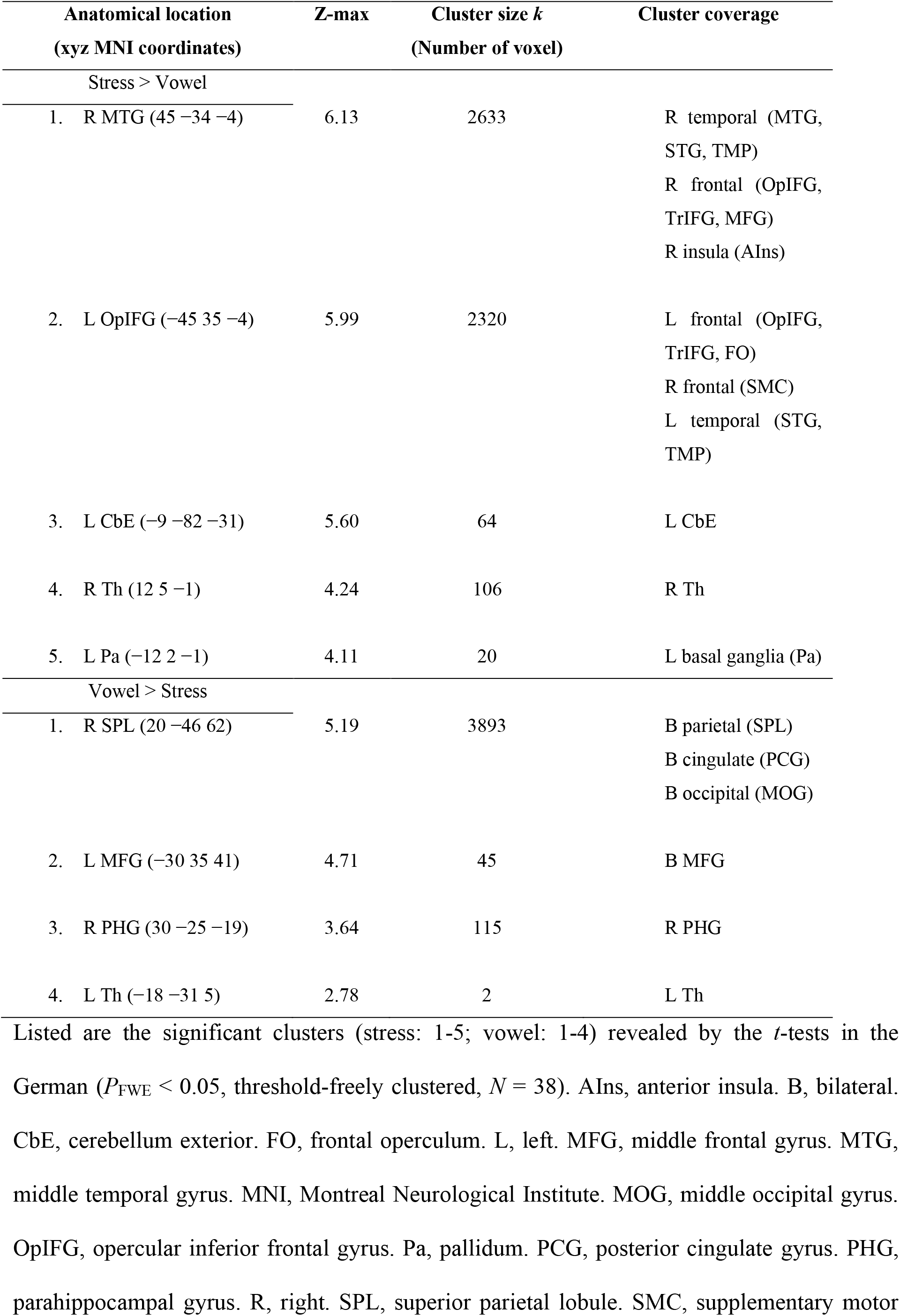
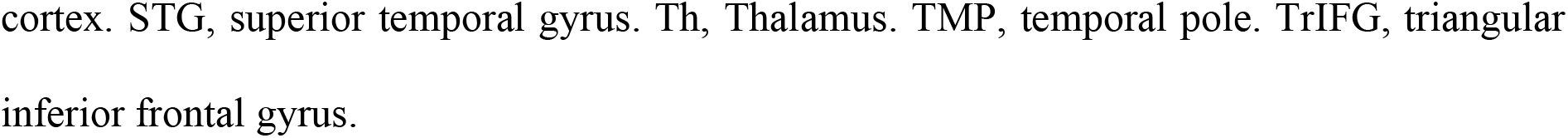
German speakers: Brain regions involved in stress and vowel processing.

**Figure 5.**
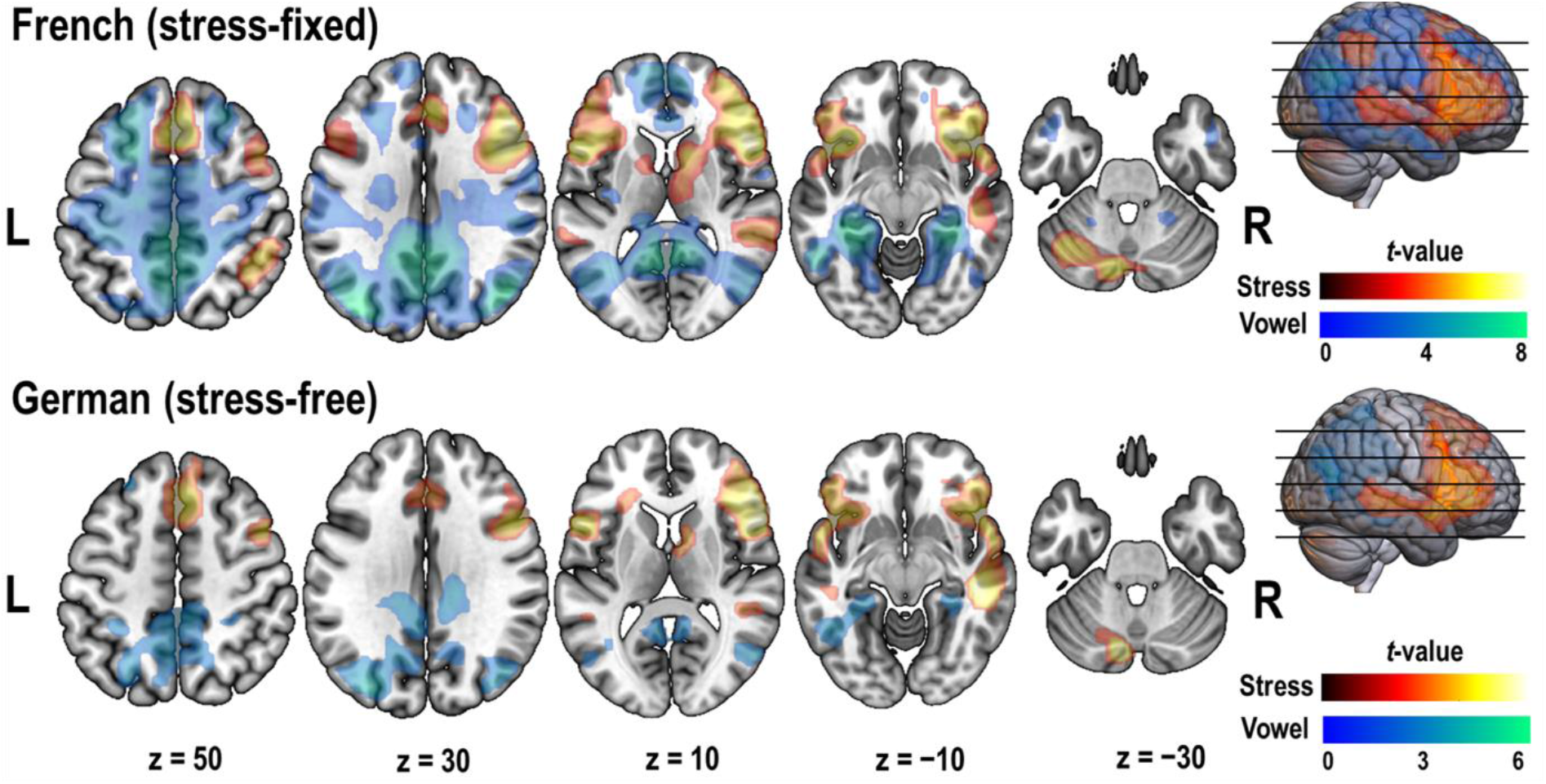
Contrast activation of both processing conditions for both groups. The significant activation (*P*_FWE_ < 0.05, threshold-freely clustered) revealed from the stress (stress > vowel) and vowel (vowel > stress) contrasts for the French speakers (top, *N* = 47) and German speakers (below, *N* = 38) is respectively mapped onto a standard brain template (MNI152). Apparent are 5 axial slices (x = 0; y = 0; z = 50, 30, 10, −10, −30). For cluster details, see Table 4 and 5. L, left. R, right.

## 4. Discussion

In the present study, we replicated the neuronal findings revealed from SP within native languages, namely the involvement of language-related brain areas, and expanded the generalizability of these findings to the more challenging and ecologically valid condition of foreign SP. Moreover, applying a cross-linguistic approach uncovered an additional mechanism beyond that of speech but yet reflecting a fundamental property of the brain. Our findings suggest that foreign SP operates upon an antagonistic architecture of two domain-general systems, regulating attentive engagement for counteracting challenges and compensating for SD. The two systems consist of the DMN and right-lateralized attention network.

### 4.1 Brain regions contributing to stress processing

The participants, regardless of the L1, processed stress less efficiently than vowel, as evident from the rise of error rates when discriminating stress patterns from pairs of stress-free foreign words. This finding is in line with past research, ascribing a strong cognitive component to SP (Dupoux et al., 2008, 2001, 1997; Heisterueber et al., 2014; Honbolygó et al., 2019; Lu et al., 2018; Schwab et al., 2020). Accordingly, our stress-contrasts suggest the involvement of the right inferior parietal lobule (i.e., AnG, SMG) and AIns as well as a bilateral involvement of frontal areas with emphasis on the IFG, MFG and SMA (Aleman et al., 2005; Domahs et al., 2013; Heisterueber et al., 2014; Kandylaki et al., 2017; Klein et al., 2011). Altogether, these structures contribute to higher cognitions not limited to speech but comprising domain-general sequence processing and salience detection as well as executive functions including working memory (Cona and Semenza, 2017; Fedorenko and Blank, 2020; Igelström and Graziano, 2017; Singh-Curry and Husain, 2009; Uddin, 2015; Uddin et al., 2017; Wager and Smith, 2003). Our stress-contrasts further highlight the bilateral involvement of auditory-processing cortical areas (i.e., STG, MTG (Domahs et al., 2013; Heisterueber et al., 2014; Honbolygó et al., 2020; Kandylaki et al., 2017; Klein et al., 2011)) and of speech-contributing subcortical structures belonging to the cerebellum, thalamus and basal ganglia (Silveri, 2021; Skipper and Lametti, 2021). By contrast, the participants required less cognitive load for processing vowel, as evident by a better performance in discriminating vowels in foreign word pairs and by the lack of activation across the fronto-insular and temporal region. Instead, our vowel-contrasts revealed activation at medial-posterior areas, suggesting the involvement of the PCG for mere attention regulation (Leech and Sharp, 2014), as well as the involvement of the PHG and the bilateral parieto-occipital region contributing to mental imagery, mnemonic and phonological processing (Cavanna and Trimble, 2006; Seghier, 2013; Ward et al., 2014).

### 4.2 Stress deafness and functional antagonism

By testing samples of both stress-familiar and “stress-deaf” participants, as confirmed by the stress-specific performance drop in discriminating pairs of stress-free foreign words, we were able to disentangle differential brain functioning contributing to SP at different proficiency levels. Furthermore, our two experimental conditions, both involving verbal tasks, allowed us to control for the contribution of language-related brain areas, shedding light on the domain-general aspect of SP (Alfano et al., 2010; Dupoux et al., 2008, 2001, 1997; Honbolygó et al., 2019; Lu et al., 2018; Peperkamp et al., 2010; Schwab et al., 2020; Schwab and Dellwo, 2017). The revealed group-condition interaction effect from our analyses suggests that SP in stress-free foreign language requires attentional resources to a degree in inverse proportion to the proficiency level, and operates antagonistically upon certain core structures that are unspecific to speech but instead belong to a right-lateralized attention system (i.e., MFG, AIns) and the DMN (i.e., PCG, MSFG, PHG). One core structure represents the right MFG, a functionally overlapping region of both the ventral/bottom-up and dorsal/top-down attention system, bearing likely an integrative function between the two (Corbetta et al., 2008; Fox et al., 2006) and thus enabling the dynamic control of attention required for the central executive, the steering component of working memory (Baddeley, 1996, 1992). Thus, the right MFG as a core structure supports the previously well-reported association between SD and working memory (Dupoux et al., 2008, 2001, 1997; Heisterueber et al., 2014; Honbolygó et al., 2019; Lu et al., 2018; Schwab et al., 2020). The other attention-related core structure represents the right AIns. The insula processes salience (Uddin, 2015) and in a linguistic context likely contributes to the detection of stressed (i.e., salient) syllables within words (Wong et al., 2004). The right insula together with the right prefrontal cortex are known to function as a switch, deactivating the DMN (Sridharan et al., 2008) when the brain engages in attentive activities. This relationship between the DMN and the attention system was found to be antagonistic (or anticorrelated) in many studies (Buckner et al., 2008; Fox et al., 2005; Raichle, 2015), likely exhibiting a specialized intrinsic organization of the brain tuned for optimized functioning (Anticevic et al., 2012). The efficiency of goal-directed processing depends not only on the appropriate recruitment of specific brain areas but also on the suppression of goal-irrelevant functions and consequently the filtering out of potentially interfering cognitions perpetuated, for example, by self-referential processing and mind-wandering. This principle underlying attentive engagement was mirrored in our findings, as SP deactivated the DMN core structures, but activated the core structures pertaining to the right-lateralized attention system. Participants with a stress-fixed L1 (i.e., French) required considerable attentional resources for processing stress in a stress-free foreign language, likely reflecting a compensatory approach of a brain not sensitized to stress features. In French speakers, this attentive engagement was reflected in terms of a strong change in the hemodynamic signal at the core structures across the attention system (i.e., activation) and the

DMN (i.e., deactivation). By contrast, participants with a stress-free L1 (i.e., German processed stress in stress-free foreign language at a higher proficiency level with far less attentional resources, possibly due to the support of available internalized stress-encoding strategies. Thus, their attentive engagement was reflected in terms of a diminished change in the hemodynamic signal across the same core structures.

### 4.3 Conclusions

In line with previous studies focusing on SP within single native languages (Domahs et al., 2013; Heisterueber et al., 2014; Honbolygó et al., 2020; Kandylaki et al., 2017; Klein et al., 2011), our findings suggest that processing stress in a stress-free foreign language recruits a bilateral widespread cerebral network together with particular insular, subcortical and cerebellar structures. However, SP was also reported to underlie a functional lateralization, with current research favoring the dominance of both either the left or right hemisphere (Aleman et al., 2005; Gandour et al., 2004; Häuser and Domahs, 2014; Klein et al., 2011; Sammler et al., 2015). In context of current neurophysiological models (Boemio et al., 2005; Bornkessel-Schlesewsky et al., 2015; Bornkessel-Schlesewsky and Schlesewsky, 2013), the process of prosodic segmentation is ascribed to the dorsal stream, the pathway responsible for time-dependent computations covering the primary auditory cortex, posterior STS, inferior parietal lobule, premotor cortex, and posterior and dorsal parts of the IFG. In particular, the processing of prosodic information within short segments, such as syllables, has predominantly been ascribed to the more temporally sensitive left hemisphere (Boemio et al., 2005). By contrast, our results provide evidence for a rightward lateralization modulating SP in a stress-free foreign language that overlaps with the location covered by the dorsal stream but remains unspecific to speech. Instead, our right-lateralized findings reflect domain-general attentive processing.

## Data availability

Datasets have been deposited in the open access repository, Zenodo (DOI: 10.5281/zenodo.7031880) (Mouthon et al., 2022). Raw MRI are organized according to the Brain Imaging Data Structure (BIDS) (Gorgolewski et al., 2016).

## Acknowledgments

We thank Profs. Nathalie Giroud (University of Zurich) and Rachel Theodore (University of Connecticut) for their advice on designing the fMRI paradigm; Laura Canedo for recording the Spanish words, and Justine Salvadori, Eugenia Ferreira da Silva and Ilona Yakoub for their help with the fMRI data collection.

